# A targeted computational screen of the SWEETLEAD database reveals FDA-approved compounds with anti-dengue viral activity

**DOI:** 10.1101/2020.07.09.194993

**Authors:** Jasmine Moshiri, David A. Constant, Bowen Liu, Roberto Mateo, Steven Kearnes, Paul Novick, Ritika Prasad, Claude Nagamine, Vijay Pande, Karla Kirkegaard

**Affiliations:** Department of Microbiology and Immunology, Stanford University School of Medicine, Stanford, California, United States of America; Department of Genetics, Stanford University School of Medicine, Stanford, California, United States of America; Department of Biology, Stanford University School of Medicine, Stanford, California, United States of America; Department of Chemistry, Stanford University School of Medicine, Stanford, California, United States of America; Department of Comparative Medicine, Stanford University School of Medicine, Stanford, California, United States of America

## Abstract

Affordable and effective antiviral therapies are needed worldwide, especially against agents such as dengue virus that are endemic in underserved regions. Many antiviral compounds have been studied in cultured cells but are unsuitable for clinical applications due to pharmacokinetic profiles, side effects, or inconsistent efficacy across dengue serotypes. Such tool compounds can, however, aid in identifying clinically useful treatments. Here, computational screening (Rapid Overlay of Chemical Structures) was used to identify entries in an *in silico* database of safe-in-human compounds (SWEETLEAD) that display high chemical similarities to known inhibitors of dengue virus. Molecules known to inhibit dengue proteinase NS2B/3, dengue capsid, and the host autophagy pathway were used as query compounds. Following computational and initial virological screening, three FDA-approved compounds that resemble the tool molecules structurally, cause little toxicity and display strong antiviral activity in cultured cells were selected for further analysis. Pyrimethamine (IC_50_ = 1.2 µM), like the dengue proteinase inhibitor ARDP0006 to which it shows structural similarity, inhibited intramolecular NS2B/3 cleavage. Lack of toxicity allowed testing in mice, in which pyrimethamine also reduced viral loads. Niclosamide (IC_50_ = 0.28 µM), like dengue core inhibitor ST-148, affected structural components of the virion and inhibited early processes during infection. Vandetanib (IC_50_ = 1.6 µM), like cellular autophagy inhibitor spautin-1, blocked viral exit from cells and could further be shown to extend survival *in vivo*. Our approach confirmed previous studies which used extensive high-throughput screening to identify niclosamide and pyrimethamine as antivirals, and it also revealed their likely molecular targets. Thus, three FDA-approved compounds with promising utility for repurposing to treat dengue virus infections and their potential mechanisms were identified using computational tools and minimal phenotypic screening.

**Author Summary:** No antiviral therapeutics are currently available for dengue virus infections. By computationally overlaying the 3D chemical structures of compounds known to inhibit dengue virus with those of compounds known to be safe in humans, we identified three FDA-approved compounds that are attractive candidates for drug repurposing towards treatment of dengue virus infections. We identified potential targets of two previously identified antiviral compounds and revealed a previously unknown potential anti-dengue drug, vandetanib. This computational approach to analyze a highly curated library of structures has the benefits of speed and cost efficiency. It also leverages mechanistic work with query compounds used in biomedical research to provide strong hypotheses for the antiviral mechanisms of the safer hit compounds. This workflow to identify compounds with known safety profiles in humans can be expanded to any biological activity for which a small-molecule query compound has been identified, potentially expediting the translation of basic research to clinical interventions.

## Introduction

Infectious disease poses a growing risk to global public health, disproportionately affecting underprivileged populations. Dengue virus is responsible for an estimated 390 million infections each year [1], yet despite this enormous global public health and economic burden [2], there are currently no approved antivirals available to treat dengue infections. The costs of research and development and the specter of drug resistance are significant challenges associated with developing novel antiviral medications to combat this disease and others caused by RNA viruses [3]. Dengue virus infects cells by receptor-mediated endocytosis of the enveloped virus. Within the endosome, acidification and ubiquitination [4] facilitate disassembly of the dengue viral capsid, an oligomer of dengue core protein, and the subsequent release of the single-stranded, positive-sense RNA genome. The viral RNA is translated in a continuous open reading frame to generate a large polyprotein that must be cleaved to free nonstructural and structural viral proteins. Dengue relies on host proteinases and its own viral proteinase, NS2B/3, to process the polyprotein. RNA replication is performed by negative-strand synthesis, followed by positive-strand synthesis, in membrane-associated complexes. Virion assembly also occurs in adjacent membranous complexes crafted from lipid droplets [5]. Virions are subsequently processed through the Golgi for maturation and released through the secretory pathway while co-opting components of the cellular autophagy pathway for viral maturation and release [6]. Throughout its infectious cycle, dengue virus relies on many other host factors to facilitate its growth as well (e.g. [7–14]).

Developing effective antivirals is especially challenging because of the propensity for drug-resistant viral variants to emerge and rapidly dominate a population. Like most RNA viruses, dengue virus has a high mutation rate and exists as a quasi-species within every infected cell [15]. This diversity can lead to the rapid emergence of drug resistance upon the application of selective pressure. Preferentially choosing compounds that suppress the rapid fixation of drug-resistant viral variants from the quasi-species can lower risk of antiviral resistance. One means by which this can be achieved by combination therapy, in which multiple compounds are used so that the probability of the presence of a drug-resistant genome is reduced. Similarly, inhibiting the pro-viral activity of a host factor could reduce the probability of drug resistance because the mutation frequency of the host is low [16]. Another approach is to accept that, for any individual compound, drug-resistant genomes will inevitably be formed during error-prone RNA replication. However, their outgrowth can be blunted if the remaining drug-susceptible genomes within the same are genetically dominant. Examples of such ‘dominant drug targets’ are viral capsids, which are formed by the intracellular oligomerization of individual capsid proteins derived from multiple genomes [17,18].

Other challenges associated with antiviral drug development may be alleviated by adopting strategic practices to reduce research and development expenses and timelines. The cost of bringing a single drug through the development pipeline, recently estimated at $2.6 billion, is extremely high and rising further [19]. One strategy to bypass costly stages of the development pipeline and reduce time-to-market is to focus on repurposing compounds that have been previously approved for other indications [20]. Through previous studies, these drugs will have undergone safety evaluations in preclinical models and human patients, decreasing the risk of failure during Phase I clinical trials or even allowing some clinical trial stages to be bypassed altogether. An additional strategy, the implementation of computational screening methods, can expedite and lower costs of preliminary high-throughput analyses compared to traditional first-round phenotypic screening.

As is the case for most RNA viruses, treatment options for dengue infections are limited to supportive therapy and do not include direct-acting antivirals [21]. However, many inhibitors of dengue virus with known targets have been reported in the literature. While these inhibitors have pharmacokinetic profiles, side effects, or inconsistent efficacy across dengue serotypes that make them unsuitable for clinical use, they are useful for investigating the underlying biology (Fig. 1A). Compounds previously investigated in this laboratory and others include: ARDP0006, which inhibits NS2B/3 proteinase activity [22], especially at one intramolecular cleavage site within NS3 [23]; ST-148, a compound that hyperstabilizes core protein interactions and has been shown to display reduced selection for resistant viruses due to the dominance of drug-susceptible genomes [24,25]; and spautin-1, an inhibitor of cellular autophagy [26] which has been shown to disrupt virion maturation [6].

**Fig 1.**
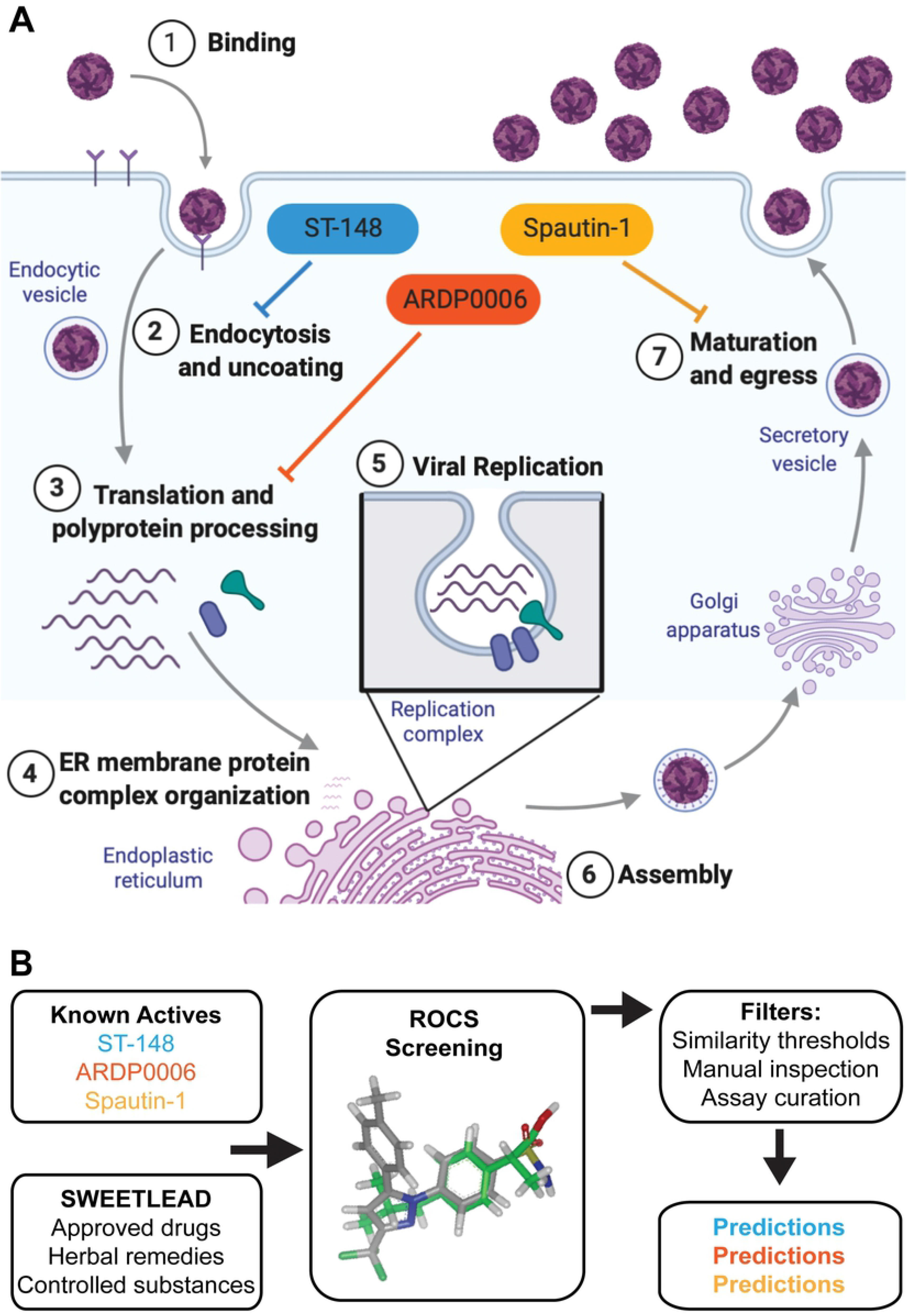
*In silico* screening to identify candidates for anti-dengue repurposing using known query compounds. (A) single infectious cycle of dengue virus is shown diagrammatically to illustrate some of the multiple steps at which antivirals can interfere. Core-binding molecules such as ST-148 can be directly virucidal, inhibit genome release (step 2), and interfere with assembly (step 6). Proteinase inhibitors such as ARDP0006 derange polyprotein processing and all subsequent stages that require mature proteins (step 3 and beyond). Inhibitors of autophagy and other membrane-sculpting processes can interfere with assembly of RNA replication complexes (step 5) or, like spautin-1, with maturation and egress (step 7). Created with BioRender.com. (B) The SWEETLEAD *in silico* database of high-confidence chemical structures for FDA-approved compounds, medicinal herbs, and controlled substances was searched to identify those with high 3D shape and chemical similarity to query compounds ARDP0006, ST-148, and spautin-1. The Rapid Overlay of Chemical Structures (ROCS) virtual screening tool scores molecular similarity between pairs of molecules based on 3D shape overlap and chemical similarity. The resulting compound hits ranked by the Tanimoto combo score were manually inspected for similar molecular features and filtered based on known side effects and commercial availability (S1 Table in supplemental material).

In this work, we aimed to identify therapeutics with potential for repurposing towards treatment of dengue virus. We used ARDP0006, ST-148, and spautin-1 as our queries to computationally search a highly curated library of safe-in-human compounds with the goal of identifying those with three-dimensional chemical similarity to our known antiviral query compounds [27]. For each query compound, we identified a safe-in-human hit that strongly inhibited dengue virus replication in tissue culture and exhibited antiviral mechanisms similar to the initial query compound. This approach offers advantages compared to traditional screens, namely the speed and cost benefits of *in silico* work and the pre-existence of strong hypotheses for mechanisms of action based on the known functions of the previously studied compounds.

## Results

### Selection of methodology, compound library, and query compounds for in silico screening

To identify pharmaceuticals whose safety profiles in humans are already known, we utilized a chemical library termed SWEETLEAD (<u>S</u>tructures of <u>W</u>ell-curated <u>E</u>xtracts, <u>E</u>xisting <u>T</u>herapies, and <u>L</u>egally regulated <u>E</u>ntities for <u>A</u>ccelerated <u>D</u>iscovery) [27]. SWEETLEAD is a highly curated *in silico* collection of thousands of FDA-approved compounds, medicinal herbs, and controlled substances with known activities. We applied a ligand-based virtual screening strategy based on the assumption that chemically and structurally similar compounds have similar biological activities [28]. In particular, we used ROCS (Rapid Overlay of Chemical Structures) software [29], an algorithm for ligand-based virtual screening that compares 3D shape and chemical similarity between pairs of molecules, to search the SWEETLEAD database for compounds with structural similarity to three inhibitors of dengue virus infection with known biochemical targets: ARDP0006, ST-148, and spautin-1 (Fig 1B).

### Utilization of ROCS-identified compounds with antiviral activity

For each query molecule, the top 500 compound hits obtained from the ROCS algorithm, as ranked by the Tanimoto combo score, were manually inspected for similar chemical and structural features and further prioritized by other factors such as a lack of known side effects and commercial availability. Several potential hits for each query compound that were reported to have limited toxicity and were not restricted by intellectual property considerations were tested for their effects on dengue virus growth and toxicity in BHK-21 and Huh7 cells (S1 Table in the supplemental material). Altogether, of the 15 candidates tested, three were selected for further study: pyrimethamine (a hit of ARDP0006), niclosamide (a hit of ST-148), and vandetanib (a hit of spautin-1).

### Pyrimethamine inhibits NS2B/3 proteinase cleavage

The NS2B/3 proteinase of dengue virus cleaves itself and other nonstructural viral proteins out of the nascent polyprotein and is necessary both for efficient polyprotein processing and cleaving host restriction factors [30,31]. In the presence of NS2B/3 proteinase inhibitor ARDP0006, we have presented evidence that inhibition of obligately intramolecular self-cleavage events results in accumulation of uncleaved or partially cleaved polyprotein precursors that are toxic to viral replication [23]. Some of this evidence includes the fact that inhibition of viral growth in cell culture by this compound (IC_50_ = 2.7 µM) [32] is much more potent than its inhibition of NS2B/3 proteinase cleavage activity in solution (IC_50_ = 620 µM), an observation we attribute to the accumulation of such toxic precursors [23]. We found that pyrimethamine, an anti-malarial FDA-approved drug, shared 3D chemical structural similarity with ARDP0006 (Fig 2A). This similarity includes the overlap of core aromatic rings of the two compounds as well as overlap between a pair of hydroxyl groups in ARDP0006 with a pair of primary amine groups on pyrimethamine (Fig 2B). Therefore, we sought to determine whether pyrimethamine inhibits dengue virus growth by a mechanism similar to that of ARDP0006.

**Fig 2.**
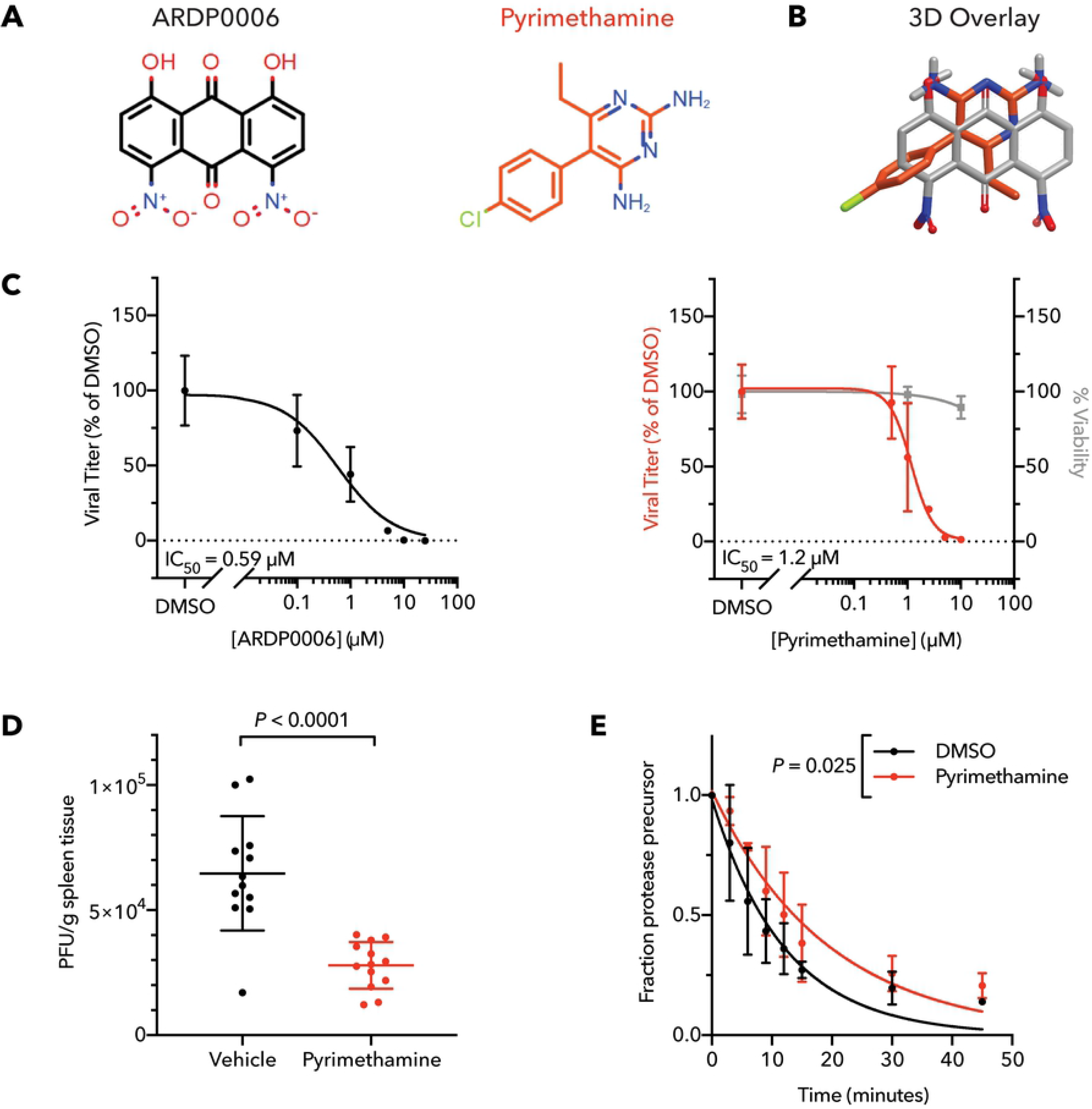
Pyrimethamine inhibits dengue virus and NS2B/3 proteinase cleavage like ARDP0006. (A) Chemical structures and (B) ROCS overlay of ARDP0006 and pyrimethamine. (C) Dose-response curves for ARDP0006 and pyrimethamine treatment of DENV2-infected Huh7 cells revealed IC_50_ values of 0.59 µM and 1.2 µM, respectively. Compounds were added 1 h prior to infection, removed, then re-administered after removal of viral inoculum. Infectious virus in cell supernatant was quantified by plaque assay at 24 hpi (left y-axis, black or red). Cell viability was measured at the same pyrimethamine doses 24 h post-treatment by WST-1 assay (right y-axis, grey). Viral titer and cell viability were normalized to DMSO-treated samples. (D) Splenic titers of DENV2-infected AGB6 mice at 4 days post-infection. Mice were treated with pyrimethamine (*N* = 13, 40 mg/kg) or vehicle control (*N* = 12) and data were analyzed by two-tailed unpaired t-test. (E) Cleavage kinetics of NS2B/3/4A in the presence of 100 µM pyrimethamine. NS2B/3/4A protein was expressed *in vitro* and radiolabeled with ^35^S-methionine, followed by treatment with 100 µM pyrimethamine or DMSO. Reaction samples were obtained over a 45 min timecourse, separated by SDS-PAGE, and fraction of uncleaved proteinase precursor remaining was determined. *N* = 3. Data were fit using single-phase exponential decay functions and *P*-value was determined by comparing the rate constant, *K*, of the models using the extra sum-of-squares F test.

To characterize pyrimethamine’s antiviral activity, we evaluated viral titer in the presence of increasing drug concentrations. The IC_50_ of viral inhibition in a single infectious cycle was 1.2 µM (Fig 2C), only two-fold higher than the IC_50_ of ARDP0006 observed here (0.59 µM, Fig 2C). Unlike ARDP0006, pyrimethamine showed little toxicity in mice (S2 Table), and thus it could be tested for dengue inhibition *in vivo*. Significant reduction in splenic titer was observed following retroorbital inoculation of virus in dengue-susceptible mice (Fig 2D).

To determine whether pyrimethamine was an inhibitor of intramolecular NS2B/3 proteinase activity like ARPD0006, we tested its effect on the cleavage of NS2B/3/4A precursor in protein translation extracts. This *in vitro* translation-based proteinase cleavage assay demonstrated that pyrimethamine significantly slowed dengue NS2B/3 cleavage but, like ARDP0006, required a dose far greater than the antiviral IC_50_ to achieve this effect (Fig 2E). The striking difference between cell culture IC_50_ and biochemical IC_50_ suggests that pyrimethamine inhibits dengue virus in a similar manner to the query compound ARDP0006, by altering the proteolytic cleavage activity of NS2B/3 in a way that results in accumulation of uncleaved viral precursors that are toxic to essential viral processes in infected cells. We conclude that pyrimethamine does not require complete efficacy of inhibiting proteolytic activity *per se*, but rather even minor inhibition of proteinase activity can lead to effective dampening of infection.

### Niclosamide augments virucidal activity of query compound ST-148

The core protein of dengue virus is an attractive target for antiviral drug development due to its complex oligomerization state, making it a candidate for dominant drug targeting [17]. The effect of query compound ST-148 is to hyperstabilize the protein-protein interactions between monomers of core protein in the viral capsid, leading to inhibition of the early stage of viral uncoating as well as later assembly steps [24,25]. We identified niclosamide, an anti-helminthic drug, as having structural similarity to ST-148 (Fig 3A). Figure 3B shows that niclosamide and ST-148 display overlap of both the central aromatic rings and the amide bonds. Additionally, there is a good overlap between the primary amine group in ST-148 and the hydroxyl group in niclosamide.

**Fig 3.**
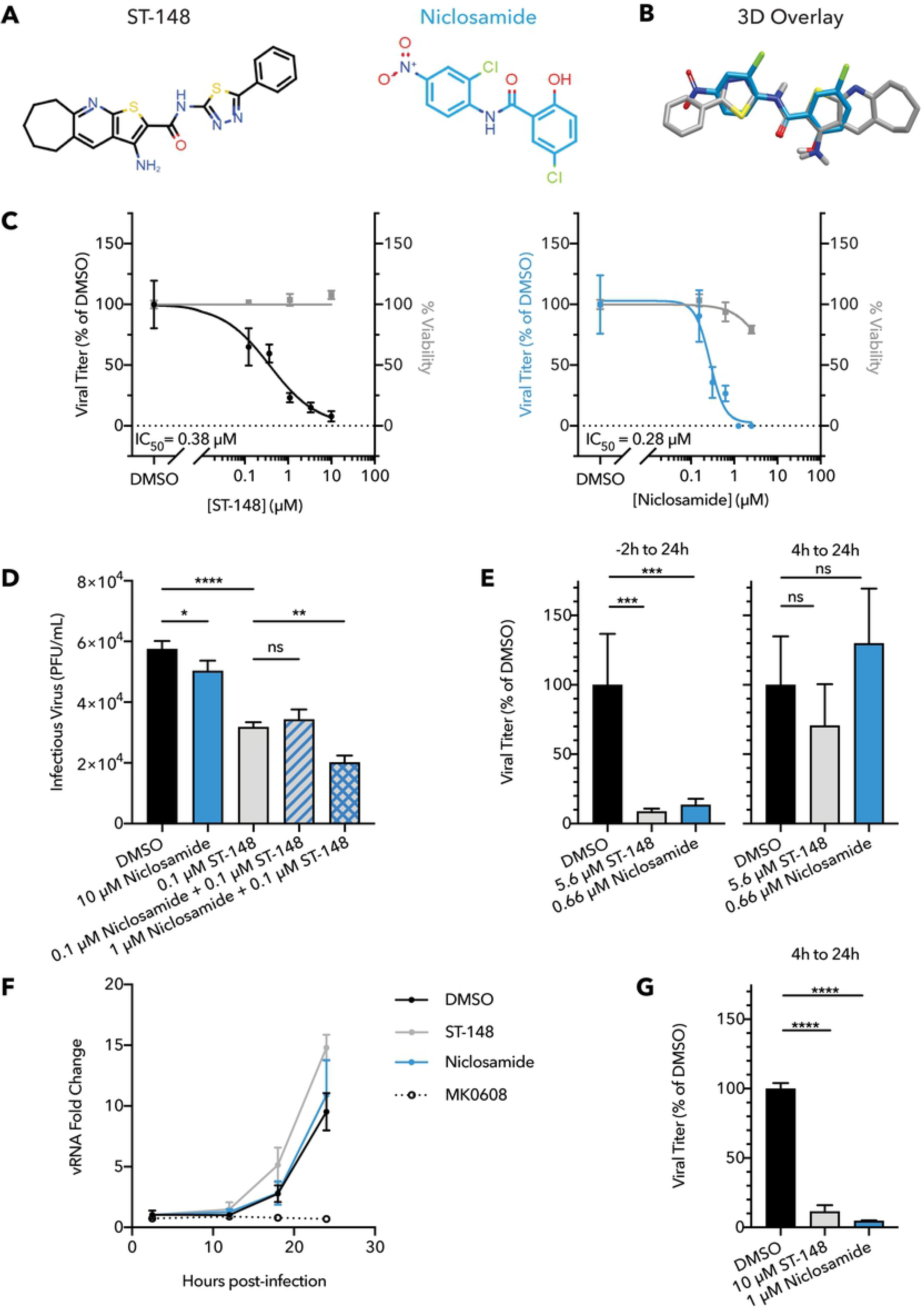
Niclosamide inhibits dengue virus at multiple stages and augments ST-148 virucidal activity. (A) Chemical structures and (B) ROCS overlay of ST-148 and niclosamide. (C) Dose-response curves for ST-148 and niclosamide treatment of DENV2-infected Huh7 cells revealed IC_50_ values of 0.38 µM and 0.28 µM, respectively. Compounds were added 1 h prior to infection, removed, then re-administered after removal of viral inoculum. Infectious virus in cell supernatant was quantified by plaque assay at 24 h post-infection (left y-axis, black or blue). Cell viability was measured at the same compound doses 24 h post-treatment by WST-1 assay (right y-axis, grey). Viral titer and cell viability were normalized to DMSO-treated samples. (D) Virucidal activity of each compound was measured after 60 min co-incubation of drugs with viral stock at 37 °C. Viral titer quantified by plaque assay on BHK-21 cells. Combination of 1 µM niclosamide + 0.1 µM ST-148 significantly increased virucidal activity over 0.1 µM ST-148 alone. *N* = 3, one-way ANOVA with Sidak’s multiple comparisons test, \**P* < 0.05, \*\**P* < 0.01, \*\*\*\**P* < 0.0001. (E) Infected Huh7 cells were treated with ST-148 and niclosamide at the IC_90_ doses for different durations within a single cycle 24 h infection. Graph titles indicate the duration of drug treatments. On the left, cells were pre-treated for 2 h prior to infection, removed during a 1 h infection at MOI = 0.5, then re-administered after removal of viral inoculum. On the right, cells were similarly infected for 1 h at MOI = 0.5 and compounds were first introduced 4 h after starting the infection. In both cases, cell supernatant was collected at 24 hpi and infectious virus was quantified by plaque assay. *N* = 4, one-way ANOVAs with Dunnett’s multiple comparisons tests, *P* > 0.05 = ns, \*\*\**P* < 0.001. (F) ST-148 and niclosamide did not reduce viral RNA replication in infected Huh7 cells. Cells were infected for 1 h with dengue virus at an MOI = 0.5, and at 4 hpi compounds were administered at the IC_90_ doses of 5.6 µM ST-148 or 0.66 µM niclosamide. Compounds were added at 4 hpi to allow viral entry. At several timepoints during the first cycle of infection, intracellular RNA was collected and quantified by qRT-PCR. Only the control for inhibited vRNA replication, 50 µM MK0608 treatment, significantly reduced vRNA load relative to the 1% DMSO vehicle control (*P* < 0.001 at 18 hpi, *P* < 0.0001 at 24 hpi). *N* = 4, two-way ANOVA with Dunnett’s multiple comparisons test. (G) When administered at an increased dose, ST-148 and niclosamide were effective during later, post-entry stages. 10 µM ST-148 or 1 µM niclosamide were administered after 4 h of infection in Huh7 cells, and extracellular virus was collected at 24 hpi and quantified by plaque assay. *N* = 3, one-way ANOVA with Dunnett’s multiple comparisons test, \*\*\*\**P* < 0.0001.

The ability of niclosamide to inhibit dengue virus infection was tested in single-cycle growth curves (Fig 3C). The IC_50_ for niclosamide inhibition of infectious extracellular virus production (0.28 µM) was comparable to that observed for ST-148 (0.38 µM). Capsid-binding viral inhibitors can be directly virucidal, inhibit viruses through cellular mechanisms such as entry and assembly, or both. To test whether ST-148 or niclosamide is directly virucidal, infectious virus stocks were incubated in the presence of drugs or DMSO controls for one hour at 37 °C, after which viral infectivity was quantified by plaque assay in the absence of compounds. We observed that co-incubation of virions with just 0.1 µM ST-148 significantly reduced viral infectivity (Fig 3D). At a much higher concentration, niclosamide was also significantly, albeit slightly, virucidal. However, addition of 1 µM niclosamide to 0.1 µM ST-148 significantly increased the virucidal effect of ST-148. Thus, we hypothesize that niclosamide binds directly to the viral core protein where it can allosterically enhance binding of ST-148. The differences in the steepness of the inhibition curves (Fig 3C) further suggests that niclosamide binds more cooperatively to virions than ST-148.

Previous reports on the inhibition of dengue virus by niclosamide have argued for vulnerable points in the viral infectious cycle: at an early step, preventing the endosomal acidification required for uncoating; at a protein processing step, by inhibiting NS2B/3 proteinase activity; and at a later step, by interfering with virion maturation [33–35]. To test a requirement for ST-148 and niclosamide at early times in infection, we compared the effects of IC_90_ doses when the compounds were present for the full timecourse of infection or administered at four hours after infection, to allow early steps of viral entry and uncoating. Delayed addition of either compound eliminated the observed reduction of viral titer (Fig 3E), indicating that, at this low concentration, the relevant inhibition occurred during the first four hours of infection. To test whether this early effect was on cell entry or any subsequent process such as translation or RNA synthesis, we tested the effect of these compounds on viral RNA abundance, which requires the accumulation of protein as well as the synthesis of RNA. When either compound was added at four hours post-infection at its IC_90_ dose, no difference in viral RNA abundance was observed, whereas the addition of MK0608, a potent inhibitor of viral RNA synthesis, strongly reduced viral RNA abundance (Fig 3F). Thus, we conclude that the most critical effects of both ST-148 and niclosamide occur early in the viral infectious cycle, during the process of receptor binding, cell entry, or genome release. Niclosamide’s antiviral activity during early infection has been previously attributed to blocking endosomal acidification; however, drug binding to core protein might affect this stage as well, as a virion with hyperstabilized core protein interactions would be incapable of uncoating. Finally, given previous literature suggesting that niclosamide can target late stages of viral replication, we increased the doses of niclosamide and ST-148 administered after the entry and uncoating steps. At the increased doses, both compounds inhibited virion production (Fig 3G), confirming that further inhibitory effects of niclosamide at later steps in the infectious cycle remain possible [35].

### Vandetanib and query compound spautin-1 disrupt viral egress independent of the autophagy-inducing VPS34 complex

We and others have observed that dengue virus growth is greatly enhanced by components of the cellular autophagy pathway [6,36,37]. Autophagy components that enhance dengue virus infection include VPS34, a kinase present in complex with Beclin-1 and needed for initiation of canonical autophagy; ATG9, which is involved in lipid acquisition; and LC3, which is needed for cargo loading and induction of membrane curvature. Other canonical autophagy factors, specifically ULK1, Beclin-1, and ATG5 are dispensable during dengue virus infection [37].

The most potent small-molecule inhibitor of the early stages of autophagy is spautin-1 (specific and potent autophagy inhibitor 1), which we have previously found to inhibit dengue virus growth by disrupting viral maturation and release [6]. When spautin-1 was used as a query compound, ROCS analysis of the SWEETLEAD database identified an FDA-approved drug, vandetanib (Fig 4A), as having structural similarity. Figure 4B shows that spautin-1 and vandetanib share a common central scaffold and that there is some overlap of the side benzene rings. In a dose-response assay, vandetanib inhibited dengue infection in Huh7 cells with an IC_50_ of 1.6 µM (Fig 4C), comparable to that of spautin-1 (1.1 µM). Furthermore, vandetanib was only mildly toxic (S2 Table), allowing *in vivo* testing in mice susceptible to dengue virus pathogenesis. Oral dosing of dengue virus-infected mice showed that vandetanib significantly extended survival following dengue infection (Fig 4D).

**Fig 4.**
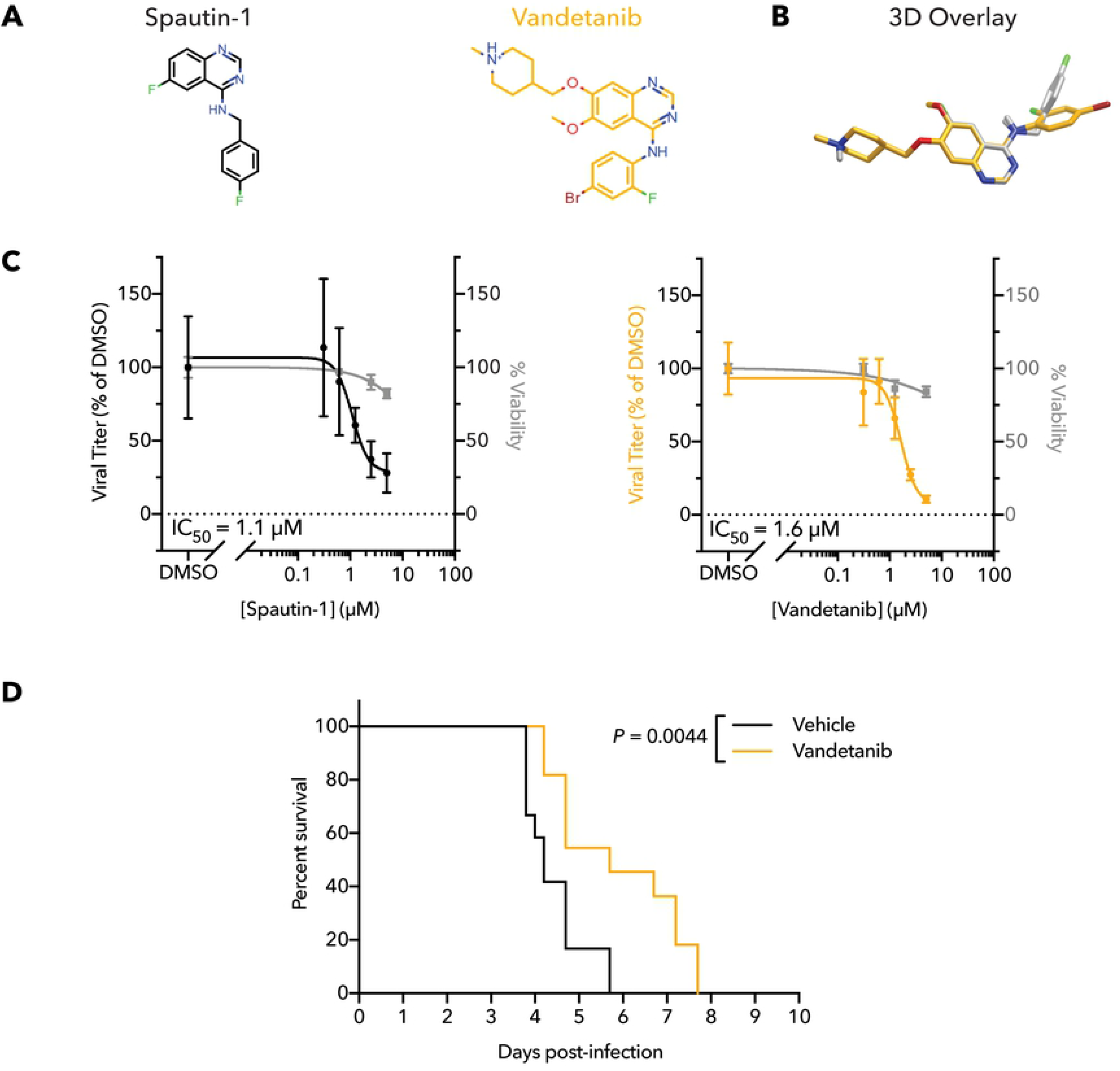
Vandetanib inhibits dengue virus *in vitro* and extends survival in infected mice. (A) Chemical structures and (B) ROCS overlay of spautin-1 and vandetanib. (C) Dose-response curves for spautin-1 and vandetanib for treatment of DENV2-infected Huh7 cells revealed IC_50_ values of 1.1 µM and 1.6 µM, respectively. Compounds were added 1 h prior to infection, removed, then re-administered after removal of viral inoculum. Infectious virus in cell supernatant was quantified by plaque assay at 24 hpi (left y-axis, black or orange). Cell viability was measured at the same compound doses 24 h post-treatment by WST-1 assay (right y-axis, grey). Viral titer and cell viability were normalized to DMSO-treated samples. *N =* 4. (D) Administration of 15 mg/kg vandetanib significantly extended survival of DENV2-infected AGB6 mice. Log-rank (Mantel-Cox) test, *P* = 0.0044. *N* = 12 (vehicle-treated) or *N* = 11 (vandetanib-treated).

To test whether vandetanib, like spautin-1, inhibited only post-RNA replication stages of dengue growth, we compared the amount of extracellular virus (Fig 5A) with intracellular virus (Fig 5B). After 24 h of infection in the presence of spautin-1 or vandetanib at their calculated IC_90_ doses (2.4 µM and 3.6 µM, respectively), virus in the extracellular supernatant was significantly reduced, as expected (Fig 5A). However, after infected cells were washed, collected, and lysed by repeated freeze/thaw, we observed that the abundance of infectious intracellular virus was unchanged by spautin or vandetanib treatment (Fig 5B). These results reveal that both spautin and vandetanib inhibit dengue virus at the stages of viral egress at this dosage. To determine whether earlier steps of viral protein synthesis and RNA amplification were truly unaffected by vandetanib, we tested the effect of the drug on the amplification of dengue virus replicon RNA. When a DENV-2 luciferase-expressing replicon was delivered by RNA transfection into Huh7 cells, dosing with vandetanib had no effect on luciferase production. This indicates that replicon RNA synthesis and protein expression were not impacted (Fig 5C), consistent with an effect of the drug at only the latest stages of viral growth.

**Fig 5.**
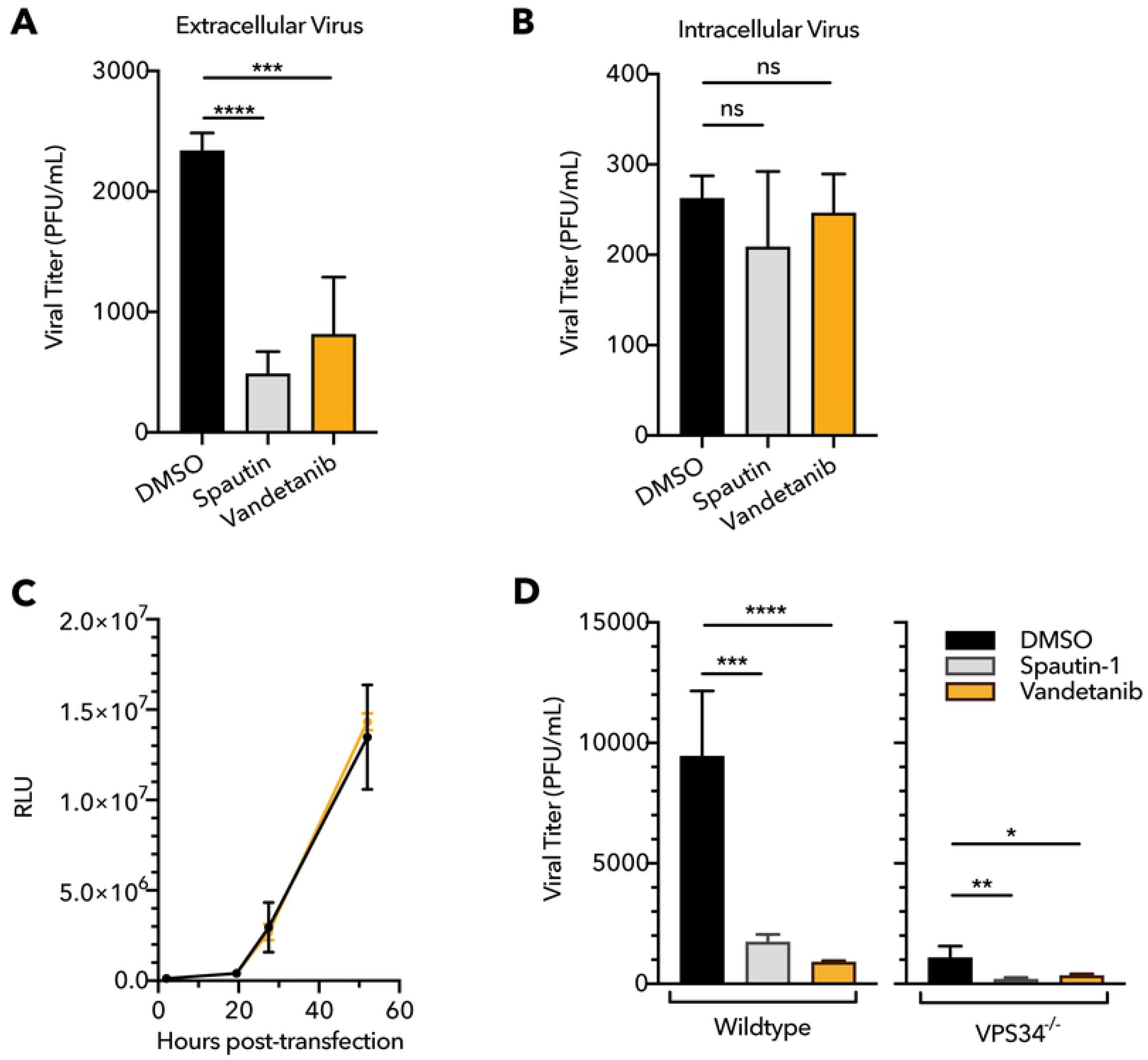
Vandetanib and spautin inhibit dengue maturation and egress in a VPS34-independent manner. Comparison of (A) extracellular and (B) intracellular viral titer at 24 hpi during treatment at the IC_90_ doses of spautin-1 (2.4 µM) or vandetanib (3.6 µM) revealed abundant intracellular virus despite reduced extracellular virus in the presence of each compound. Cells were pre-treated for 1 h, infected for 1 h at MOI = 0.5, and post-treated for an additional 23 h. *N* = 4, one-way ANOVA with Dunnett’s multiple comparisons test; *P* > 0.05 = ns, \*\**P* < 0.01, \*\*\*\**P* < 0.0001. (C) Vandetanib did not impact the stage of viral RNA replication and polyprotein translation, as evaluated by luciferase from a DENV2 reporter replicon. Huh7 cells transfected with a luciferase-expressing replicon were treated with 3 µM vandetanib (orange) or 1% DMSO vehicle control (black) for 1 h prior to transfection and up to 52 h post-transfection. Luciferase intensity was quantified as a measure of viral protein abundance. *N* = 4, two-way ANOVA with Sikak’s multiple comparisons test. Statistical significance defined as *P* < 0.05; no significant differences observed between DMSO and vandetanib treatments at any timepoint. (D) VPS34 was not required for spautin or vandetanib anti-dengue activity. Wildtype and VPS34 knockout Huh7 cells were treated with 5 µM spautin, 5 µM vandetanib, or DMSO. Cells were pre-treated, infected at MOI = 0.5, and post-treated as in (A and B). At 24 hpi, cellular supernatant was collected and titered by plaque assay. *N ≥* 3. One-way ANOVA with Dunnett’s multiple comparisons test; \**P* < 0.05, \*\**P* < 0.01, \*\*\**P* < 0.001, **** *P* < 0.0001.

Given the findings that spautin-1 represses autophagy by causing degradation of the VPS34/Beclin-1 complex [26] and that VPS34 is required for efficient virus production while other components of this complex are not [37], we hypothesized that spautin-1 and vandetanib both required VPS34 to inhibit dengue virus via the autophagy pathway. To investigate this, we compared antiviral activity of spautin and vandetanib in wild-type Huh7 cells and CRISPR/Cas9-generated VPS34 knockout Huh7 cells (Fig 5D). Absence of VPS34 significantly reduced overall viral titer in DMSO-treated cells, as expected.

However, both drugs retained significant antiviral activity in the VPS34 knockout cells, indicating the molecular mechanism of these compounds is not exclusively mediated by VPS34. These results argue that both spautin-1 and vandetanib inhibit dengue egress by an unknown mechanism that may differ from the effects of spautin-1 on autophagy.

## Discussion

With no antiviral treatments currently available for patients suffering from dengue infection, there is a great unmet need for any therapeutic targeting this disease. Given that the cost of drug discovery continues to increase [19], efficient approaches to develop such therapeutics are urgently needed. In this study, we used a ligand-based virtual screening tool to compare the three-dimensional chemical similarity of anti-dengue research compounds with safe-in-human drugs in the SWEETLEAD database and identified three FDA-approved drugs from this database: pyrimethamine, niclosamide, and vandetanib.

Pyrimethamine, a folic acid antagonist used against apicomplexan parasitic infections, shares structural similarity to query compound ARDP0006, an inhibitor of the dengue NS2B/3 proteinase. Pyrimethamine reduced dengue titer with an IC_50_ of 1.2 µM and significantly reduced dengue splenic burden in infected mice. Our results are in agreement with those of *Barrows et al.,* whose authors identified pyrimethamine as an inhibitor of Zika virus, another member of the flavivirus family, in an extensive phenotypic screen of FDA-approved compounds [38]. We found that, like query compound ARDP0006, pyrimethamine affected the kinetics of intramolecular NS2B/3 cleavage, slowing proteinase cleavage *in vitro*. These results suggest that pyrimethamine inhibits dengue virus, and likely Zika, through inhibition of the NS2B/3 viral proteinase. Our previous work with ARDP0006 argued that the strong inhibition of viral growth by the relatively modest NS2B/3 proteinase inhibition in solution results from the accumulation of improperly cleaved viral products [23].

We found that niclosamide, an FDA-approved drug used to treat tapeworm infections, significantly inhibited dengue virus with an IC_50_ of 0.28 µM. This drug has been reported to have many potential therapeutic effects, including inhibition of several bacterial and viral infections, cancer, and metabolic disease, among others [38–46]. However, the direct binding targets of niclosamide for the majority of these indications remain unknown. In this work, we found that niclosamide augmented virucidal activity of ST-148, the core-targeting query compound with which it shares structural similarity. These results suggest a novel mechanism of niclosamide inhibition of dengue virus by direct interaction with the dengue core protein. Other mechanisms of viral inhibition by niclosamide, such as previously identified inhibition of NS2B/3 proteinase [33] and blocking endosomal acidification leading to ineffective entry and maturation [34,35] may also be relevant. The presence of these multiple mechanisms of anti-dengue activity, possibly targeting multiple viral and host proteins, may engender a high genetic barrier to antiviral resistance against niclosamide.

Lastly, we demonstrated anti-flaviviral activity for vandetanib, a multiple-receptor tyrosine kinase inhibitor approved for use against medullary thyroid cancer. Vandetanib inhibited dengue virus in cultured cells with an IC_50_ of 1.6 µM and significantly extended survival in a mouse model of infection. Interestingly, we observed that, at the IC_90_ concentration for extracellular virus secretion, the quantity of infectious intracellular virus was not significantly affected by drug treatment, suggesting that both vandetanib and query compound spautin-1 inhibit dengue virus at the stage of viral egress. Previous studies found that in addition to blocking egress, higher concentrations of spautin-1 also inhibit maturation of virion particles [6]. Spautin-1 has been implicated in destabilizing the Beclin-1/VPS34 complex, thus inhibiting autophagosomal initiation and maturation [26]. However, even though VPS34 is required for efficient growth of dengue virus [37], we discovered that both vandetanib and spautin-1 did not require VPS34 to limit dengue virus egress.

Vandetanib’s inhibition of viral egress could be related to its known ability to inhibit receptor tyrosine kinases such as epidermal growth factor receptor (EGFR), RET, and vascular endothelial growth factor receptor (VEGFR), which is why it is FDA-approved to treat certain cancers. Although query compound spautin-1 is best known for its stabilization of the Beclin-1/VPS34 complex via inhibition of deubiquitination enzymes USP10 and USP13, it has also been shown, like vandetanib, to inhibit EGFR signaling [47]. EGFR and other receptor tyrosine kinases are known to modulate intracellular vesicle trafficking. In fact, anti-cancer kinase inhibitors erlotinib and sunitinib exert broadly antiviral effects via this method, though unlike vandetanib treatment, these compounds reduced viral entry [48].

It is possible that vandetanib inhibition of VEGFR *in vivo* may also alleviate some of the pathology of dengue hemorrhagic fever and dengue shock syndrome because VEGFR signaling directly induces vasodilation [49]. During severe dengue, patients experience increased vascular permeability, plasma leakage, and reduced platelet count [50–52]. VEGF levels are significantly elevated in plasma from patients with severe dengue fever compared to patients with uncomplicated dengue or healthy controls [53,54]. Therefore, vandetanib inhibition of VEGFR-induced vasodilation may alleviate hemorrhaging symptoms caused by severe dengue. The possibility of directly inhibiting viral egress and simultaneously alleviating pathology of severe dengue makes vandetanib an exciting candidate for translation into a clinical setting.

When it comes to the development of antiviral drug resistance, not all targets are equal. By strategically selecting candidates with higher genetic barriers to resistance, we can develop antivirals that are more effective and have longer durations of clinical utility. As a host-targeting receptor tyrosine kinase inhibitor, vandetanib likely inhibits dengue by restricting access to a crucial host pathway, with the resulting outcome that antiviral resistance is unlikely to be rapidly selected among an existing virus population [16,55]. The target of query compound ST-148 and, most likely, its hit compound niclosamide is core protein, a viral protein for which drug susceptibility has been shown to be genetically dominant [56]. Specifically, a single monomer of dengue core protein must assemble with other core molecules to form a larger oligomer, the capsid shell. When, in the presence of drug, a single viral genome within the cell randomly arises which is drug-resistant, the monomer produced will be incorporated into a larger oligomer also containing drug-susceptible molecules, rendering the entire chimera drug-susceptible. The target of query compound ARDP0006 and hit compound pyrimethamine is NS2B/3, which is likely to be a dominant drug target as well. This proteinase must first perform multiple intramolecular proximal cleavages in *cis* to free itself from the larger polyprotein before it can cleave at more distal viral junctions. Because these cleavages occur only in *cis*, a drug-resistant proteinase would not be able to rescue a drug-susceptible neighbor in *trans*, leading to accumulation of uncleaved precursors that interfere with outgrowth of all viruses in the cell, even those with drug-resistant genomes [23]. Thus, for all three anti-dengue compounds described here, the outlook is promising in that each functions through targets for which the development of drug resistance is disfavored, which is how the query compounds were selected.

Tool compounds that are unsuitable for use in humans have been identified for many infectious agents. Structure-based *in silico* screening is emerging as a powerful technique to quickly identify candidates for drug development for many indications, including viral diseases [57–59]. The workflow used here can be easily adapted for a wide range of biological indications for which a query compound has been identified. The SWEETLEAD database of safe-in-human compounds is freely available to download in a variety of formats [27] and contains high-confidence, curated structural information for 4442 approved drugs, controlled substances, and herbal isolates. Once downloaded, the SWEETLEAD database can be searched using any computational screening workflow desired, including the ROCS virtual screening software utilized in this study [29]. Utilizing these computational tools to conduct *in silico* primary screening drastically reduced the number of compounds we were required to test in tissue culture, expediting the process of identifying candidates for repurposing and providing strong, testable hypotheses about their mechanisms. As FDA-approved drugs with antiviral mechanisms that likely disfavor drug resistance, pyrimethamine, niclosamide, and vandetanib are all promising candidates for repurposing against dengue virus that should be further investigated to meet this pressing global public health need.

## Acknowledgements

We would like to thank Jeffery Glenn, Chaitan Khosla and Peter Kim for their leadership in facilitating this collaborative work. We are grateful to Mark Smith for assistance with synthesis and acquisition of chemical compounds and to Lisa Selzer for scientific advice. We greatly appreciate critical reviews of the manuscript provided by Peter Sarnow, Molly Braun, and Lisa Selzer. We are also grateful to Daria Mochly-Rosen and Kevin Grimes, and the SPARK at Stanford program in translational research, for scientific and professional advice.

## Material and Methods

### Computational screening

Three query molecules were used for *in-silico* ligand-based screening: ST-148, ARDP0006, and Spautin-1. 3D conformers of the molecules were generated using OMEGA (version 2.5.1.4, OpenEye Scientific Software, Santa Fe, NM. http://www.eyesopen.com) [60], then compared with the chemical structures in the SWEETLEAD database (https://simtk.org/projects/sweetlead) [27], using ROCS (version 3.2.1.4, OpenEye Scientific Software, Santa Fe, NM. http://www.eyesopen.com) [29]. The top 500 hits for each query were ranked by Tanimoto combo score, which summarizes both the shape and chemical similarity, and manually inspected. Six compounds were selected for ARDP0006, seven compounds were selected for ST-148, and two compounds were selected for Spautin-1. Compounds were purchased from chemical vendors (AK Scientific, Santa Cruz Biotech, Sigma Aldrich) for experimental validation.

### Viruses and cell culture

Dengue virus strains 16681 (used in cell culture, gift of Richard Kinney) and PL046-2M (used in mouse experiments, gift of Eva Harris) [61] were produced as previously described [23]. BHK-21 cells were grown at 37 °C with 5% CO_2_ in DMEM (HyClone, GE Healthcare life sciences) supplemented with 10% bovine serum and 1 U/mL penicillin/streptomycin. Huh7 cells were grown at 37 °C with 5% CO_2_ in DMEM supplemented with 10% fetal bovine serum, 0.1 mM non-essential amino acids (Gibco), 1 mM sodium pyruvate (Gibco), and 1 U/mL penicillin/streptomycin (Gibco). Huh7 CRISPR/Cas9 VPS34 knockout cells were generated as previously described [37] and validated by Sanger sequencing and immunoblotting (S2 Fig in the Supplemental Information).

### Plaque assays

Dengue plaque assays were carried out on BHK-21 cells in 24-well plates. Cells were infected with ten-fold serial dilutions of virus, which were allowed to adsorb for at least 1 h at 37 °C, and overlaid with standard cell culture media plus 0.37% (w/vol) sodium bicarbonate and 0.8% (w/vol) Aquacide II (Millipore Calbiochem). After 7 days incubation at 37 °C, 5% CO_2_, cells were fixed with 5% (final) formaldehyde and stained with crystal violet for plaque enumeration.

### Assays for antiviral activity

Between 4×10^4^ and 1×10^5^ Huh7 cells per well were seeded into 24-well plates one day prior to infection. Compounds were diluted to 100x final concentration in DMSO, then further diluted 1:100 into appropriate cell culture. Unless otherwise indicated, tested compounds were present for 1 h pre-treatment and re-administered for post-treatment following infection. Specifically, cells were pretreated with compounds or 1% DMSO for 1 h prior to infection. Media and drugs were removed from cells for infection at MOI = 0.1 or 0.5 in 200 µL cold media per well. After 1 h at 37 °C, virus inoculum was removed by aspiration and replaced with media plus compound or control. After 24 h total infection, infectious virus was quantified by plaque assay. Anti-viral IC_50_ dose-response curves were fitted in Graphpad Prism 8 using the *[inhibitor] vs. response, variable slope* function with a constraint *bottom* > *0*. IC_90_ values are defined as the dose at which *y* = *(span* × *0.1)* + *bottom.*

### Cell viability assays

Mock-infected wells were used to determine cell viability in the presence of chemical compounds compared to 1% DMSO vehicle control. Cell viability was determined using the WST-1 kit (Abcam) per manufacturer’s instructions. The 50% cytotoxic concentration (or CC_50_) was calculated by fitting dose-response curves in Graphpad Prism 8 using the *[inhibitor] vs. normalized response, variable slope* function.

### Virucidal activity assays

Compounds were diluted to 100x in DMSO, then further diluted 1:100 into PBS containing dengue virus to a final volume of 200 µL and mixed. Aliquots were taken for plaque assays immediately after addition of compounds (T = 0 min) or after incubation at 37 °C for 60 min (T = 60 min).

### Intracellular viral RNA isolation and quantification

Supernatants were removed from infected cells and cells were washed with high salt buffer (1 M NaCl, 50 mM sodium bicarbonate, pH 9.5) for 3 min 4 °C to remove cell-associated virus [62]. RNA was extracted with TRIzol reagent (Thermo Fisher), precipitated with isopropanol, washed with 75% (vol/vol) ethanol, and resuspended in water. Positive sense vRNA and GAPDH transcripts were measured by qRT-PCR using the QuantiTect SYBR Green RT-PCR kit (Qiagen) on an Applied Biosystems 7300 machine. Fold change in dengue viral RNA was calculated using the 2^(-ΔΔC_t_) method and are relative to the early timepoint 1% DMSO control. Primer sequences used for GAPDH and viral genomic RNA are as following, in the 5’ to 3’ orientation.

GAPDH forward: CTGAGAACGGGAAGCTTGT.

GAPDH reverse: GGGTGCTAAGCAGTTGGT.

DENV NS3A forward: AATGGGTCTCGGGAAAGGAT.

DENV NS3A reverse: AAGAGCTGCTGTGAGAGTTA.

### Proteinase cleavage assays

Proteinase cleavage assays were performed as previously described [23]. Briefly, dengue protein NS2B/3/4A constructs were expressed in rabbit reticulocyte lysate (T_N_T coupled T7, Promega Bio Systems) programmed with 1 µg DNA/50 µL and labeled with 20 µCi of L-[^35^S]-methionine (EasyTag, Perkin Elmer). Reactions were incubated at 30 °C for 30 min before adding L-methionine (1 mM final) and DMSO (2% final) or pyrimethamine in DMSO. At the indicated times, 3 µL aliquots were immediately diluted with Laemmli sample buffer (4 volumes, 1X final), denatured (60 °C, 10 min.), and separated by SDS-PAGE. Gels were dried under vacuum (80 °C, 120 min.) and exposed to a low-energy phosphor storage screen (Molecular Probes). Protein quantification was performed using ImageQuant TL 8.1 software (GE Healthcare Life Sciences).

### Replicon luciferase assays

A dengue replicon derived from the dengue virus strain 16681 expressing *Renilla* luciferase and dengue nonstructural proteins has been previously established [7,63]. Huh7 cells were pretreated with compounds of interest or 1% DMSO in cell culture media for 1 h. After pre-treatment, 1 µg replicon RNA per well was transfected using Lipofectamine 3000 (Thermo Fisher Scientific) for 2 h at 37 °C in opti-MEM (Gibco). Cells were washed 3x after transfection with media, then post-treated with media plus compounds or 1% DMSO at 37 °C. Transfected cells were harvested and analyzed using the *Renilla* Luciferase Assay System (Promega BioSystems). Luciferase intensity was measured with a GloMax luminometer (10-s signal integration; Promega BioSystems).

### Intracellular virus quantification

Huh7 cells seeded into 24-well plates were pretreated with spautin-1 or vandetanib at their IC_90_ doses (2.4 µM and 3.6 µM, respectively) for 1 h at 37 °C, and infected with dengue virus at MOI = 0.5. After 1 h, virus inoculum was aspirated and replaced with media containing compounds or 1% DMSO. After 24 h total infection, extracellular virus was collected in cellular supernatant and quantified by plaque assay. To collect intracellular virus, cells were washed with PBS, incubated for 3 minutes at 4 °C with high-salt buffer (1 M NaCl, 50 mM sodium bicarbonate, pH 9.5) to detach cell-associated extracellular virus [62], washed twice with PBS, detached with 0.25% trypsin-EDTA, and isolated by centrifugation (500 ×*g*, 5 min, 4 °C). Infected cell pellets were resuspended in 750 µL cell culture media, lysed by repeated freeze/thaw in liquid nitrogen followed by 37 °C water bath. Debris was removed by centrifugation (3,200 ×*g*, 5 min, 4 °C) and infectious intracellular virus was quantified by BHK-21 plaque assay.

### Mice

Based on the original mouse models of dengue virus infection in mice lacking type I and type II interferon signaling [64–66], C57BL/6 congenic mice lacking type I and type II interferon receptors (AGB6; C57BL/6.129-*Ifnar1*^tm1Agt^ *Ifngr1*^tm1Agt^) [56] were inoculated IV with dengue virus in 100 µL PBS plus 20% FBS at 8-12 weeks of age by retro-orbital injection under ABSL2 conditions. Antiviral compound treatments were delivered by oral gavage twice daily and were begun 8-12 hours prior to infection. Pyrimethamine was prepared in PBS plus 5% v/v DMSO and 20% w/v (2-hydroxypropyl)-beta-cyclodextrin. Mice were euthanized at 4 days post-infection for determination of dengue virus quantities in splenic tissue by plaque assay. Mice received 40 mg pyrimethamine per kilogram weight delivered twice daily. Mice were housed in an AAALAC-accredited mouse facility. Husbandry is performed in accordance with the Guide for the Care and Use of Laboratory Animals, 8^th^ edition [67]. Room conditions included a temperature of 23 °C, relative humidity of 30% to 40%, and a 12:12-h light:dark cycle (lights on, 0700 h). Vandetanib was prepared in saline plus 2% v/v DMSO, 20% w/v (2-hydroxypropyl)-beta-cyclodextrin in PBS. Infected mice received 15 mg vandetanib per kilogram weight, twice daily. All experiments involving mice were approved by the Institutional Animal Care and Use Committee of Stanford University under protocol APLAC-9296.

## Supporting Information

**S1 Table. Summary of hits for lead compounds ARDP0006 and ST-148 screened for anti-dengue activity in tissue culture cells.** Hit compounds were tested for anti-dengue activity and cell toxicity in tissue culture cell lines. Summary of data depicted graphically in Fig. S1 (eliminated compounds), Fig. 2C (pyrimethamine), Fig. 3C (niclosamide), and Fig. 4C (vandetanib). Antiviral activity at the dose with the greatest effect is represented. At the compound dose with the greatest antiviral effect, not antiviral (−) defined as viral titer greater than 50% of vehicle, weakly antiviral (+) as titer between 20-50% of vehicle, moderately antiviral (++) as titer between 5-20% of vehicle, and highly antiviral (+++) as titer under 5% of vehicle. Similarly, toxicity at the compound dose with the greatest cytotoxic effect is represented. At the compound dose with the greatest cytotoxic effect, nontoxic (−) defined as cell viability greater than 80% of vehicle, mild toxicity (+) as cell viability between 60-80% of vehicle, and high toxicity (+++) as cell viability under 60% of vehicle. Two potential hits for spautin-1 were tested directly in a murine model before testing in tissue culture. One of these two hits, lapatinib, was not effective in mice and so was discontinued from further study.

**S2 Table. Acute toxicity of computationally identified, FDA-approved hit compounds.** Toxicity for hits identified in the SWEETLEAD database as having chemical similarity to query molecules as reported in the PubChem database. Data for oral and intraperitoneal administration are reported, where available. LD50 values represent acute toxicity, for a single dose by the relevant delivery route.

---

**S1 Fig. Eliminated hits for lead compound ARDP0006 and ST-148 were screened for anti-dengue activity in tissue culture cells.** BHK-21 were pre-treated with the compounds at the indicated concentrations for 30 min, infected with dengue virus at an MOI = 0.1 for 1 h, then post-treated with compound. At 48 hpi, cellular supernatant was collected and titered by plaque assay (purple). Cell viability in uninfected cells in the presence of compounds for 48 h was measured by WST assay (grey). Both viral titer and cell viability are normalized to their relative DMSO vehicle control, *N* = 2.

**S2 Fig. Validation of CRISPR-Cas9 VPS34 knockout (KO) Huh7 cells by immunoblot and Sanger sequencing.** VPS34 was knocked out in a Huh7 background through insertion of 4 bp in Exon 3, as described previously *(1)*. (A) Protein lysates from wildtype Huh7 and two separate VPS34 KO clones (clones 5 and 6) were obtained, separated by SDS-PAGE, and immunoblotted using antibodies targeted to VPS34 and GAPDH. Efficient knockout of VPS34 protein expression was observed in clone 6. (B) Genomic DNA from VPS34 clone 6 KO cells was isolated, the VPS34 region was PCR amplified around the targeted cut site, and the PCR product was sequenced by Sanger sequencing to ensure incorporation of the 4 bp insertion (highlighted in blue) by the guide RNAs. VPS34 knockout of clone 6 was determined to be successful and this clone was utilized in further experiments.

